# Plyometric training enhances strength and precision of the finger movements in pianists

**DOI:** 10.1101/2022.05.16.492083

**Authors:** Kaito Muramatsu, Takanori Oku, Shinichi Furuya

## Abstract

Stability of timing and force production in repetitive movements characterizes skillful motor behaviors such as surgery and playing musical instruments. However, even trained individuals such as musicians undergo further extensive training for the improvement of these skills. Previous studies that investigated the lower extremity movements such as jumping and sprinting demonstrated enhancement of the maximum force and rate of force development immediately after the plyometric exercises. However, it remains unknown whether the plyometric exercises enhance the stability of timing and force production of the dexterous finger movements in trained individuals. Here we address this issue by examining the effects of plyometric-like training specialized for finger movements on piano performance by well-trained pianists. The training demonstrated a decrease of the variation in timing and velocity of successive keystrokes, along with a concomitant increase in the rate of force development of the four fingers, but not the thumb, although there was no change in the finger muscular activities through the training. By contrast, such a training effect was not evident following a conventional repetitive piano practice. In addition, a significant increase in the forearm muscle temperature was observed specifically through performing the plyometric exercise with the fingers, implying its association with improved performance. These results indicate effectiveness of the plyometric exercises for improvement of strength, precision, and physiological efficiency of the finger movements even in expert pianists, which implicates a role of ways of practicing in enhancing experts’ expertise.

## INTRODUCTION

Musical performance represents one of the most skillful motor behaviors, which typically requires years of extensive musical training from childhood^1–3^. Conventional musical education and training, however, may emphasize the importance of quantity of the practice^4^ and subjective experience of trained teachers and performers, due to a lack of evidence proving effectiveness of individual ways of musical practicing^5^. In contrast, most of training and education in sports are built upon accumulated evidence through the development of sports science, which has contributed to breaking records over decades^6–8^. Following a similar perspective, musical performance requires reproducible and quantitative knowledge on the effectiveness of music education and training specialized for musicians who are required to perform highly dexterous sensorimotor skills in no way inferior to athletes.

One approach to discover the optimal way of practicing is to compare effects of different ways of practicing on the sensorimotor skills. For example, a previous study examined effects of variation of the temporal structure of piano practicing on neuromuscular control of the sequential finger movements in pianists^9^. While rhythmic variation of successive piano keystrokes in practicing improved maximum rate of keystrokes and altered finger muscular activation patterns in piano playing, there was no change in the rhythmic accuracy of the keystrokes following such a differential learning. Non-invasive brain stimulation using the transcranial direct current stimulation also improved fine control of the finger movements in untrained individuals, but not in trained pianists^10^. These results highlight difficulty of improving precision of repetitive finger movements in trained pianists, although a recent study discovered a rare case of achieving it through a specialized somatosensory training with a haptic device^11^.

Plyometric exercise has been known as one established training in the field of sports, which consists of a quick succession of eccentric and concentric contractions of the targeted muscle.^12^ Previous studies investigating this exercise have focused mainly on fast, powerful movements of the lower extremities, such as sprinting^13^ and vertical jumping^14^, and have revealed significant reductions of muscular fatigue due to a decrease in the duration of forceful contraction compared to resilience exercises that maximally stretch the muscle spindles to the same degree^12^. Post-activation performance enhancement (PAPE) has been proposed as a putative physiological mechanism underlying the short-term improvement of the performance due to an increase in rate of force development (RFD) following some physical training such as not only high-intensity resilience exercises but also plyometric training.^15^ However, evidence for the effectiveness of the plyometric exercises has been limited primarily to the lower extremity, with only a few studies in the upper extremity such as the shoulder,^16^ but none in the forearm and hand that are different from the lower extremity in terms of neurophysiological and biomechanical architectures. Also, it has not been known whether the plyometric exercises enhance fine motor control of trained individuals such as musicians.

The goal of the present study is to address effects of plyometric exercises on dexterous finger movements while trained pianists play the piano. To this aim, we assessed the time-varying trajectory of the vertical position of the piano keys, key-depression force, and finger muscular activities during the repetitive keypresses before and after the training, based on previous findings of the relationship of pianistic skills with force exertion patterns^17,18^ and muscular activities^19^. Since it has been pointed out that changes in performance due to PAPE are supported mainly by elevation of muscle temperature^20^, and since its time course has been shown to accompany changes in motor skill^21,22^, muscle temperature was measured throughout the course of time before, during, and after the training in this study. While several studies have investigated physiological mechanisms of piano performance and practicing ^19,23,24^, there has been no study assessing the skin and muscular temperature of the finger muscles during piano practicing. The present research will therefore provide performers and instructors with a basis for the application of evidence-based practice methods and training regimes, which are particularly important for the prevention of the development of overuse syndromes and focal dystonia^25^.

## METHODS

### Participants

Twenty-six pianists participated in the experiment (Nineteen females; 18-30 yr old). All of them had undergone intensive piano training and formal musical education at music conservatories and/or privately for > 14yr. The pianists were randomly classified into two groups undergoing different training tasks (see details in *Experimental Tasks*). In accord with the Declaration of Helsinki, the experimental procedures were explained to all participants. All procedures were approved by the ethics committee at the Sony Corporation.

### Experimental Setup

A digital piano with a real key action (KAWAI, VPC-1) was used in the experiment to collect data representing the timing, pitch, and velocity of the individual key presses and releases (i.e. MIDI information) with a custom-made LabVIEW (National Instruments) program. The instrumental sound was elicited via a headphone attached on the participant’s ears. The surface electromyography (EMG) system with two sets of wirelessly connected electrodes (Trigno Quattro sensors, Delsys Inc.) was connected to a laptop through an analog-to-digital board (NI USB-6363; National Instruments). Each electrode was placed on the muscle belly of the extensor digitorum communis (EDC) and flexor digitorum superficialis (FDS) of the right hand. The EMG signals were amplified, band-pass filtered (10-500Hz), and sampled at 1kHz using LabVIEW. As with EMG, a custom-made force sensor connected to an analog-to-digital convertor was used to measure the force when each finger was pressing down the sensor. A high resolution position sensor system was mounted on the bottom of the key-bed^26^, and the vertical position of the keys was recorded by 1kHz in synchronization with MIDI and EMG. Both muscle temperature of the EDC and nearby skin temperature were measured at each time point throughout the experiment (see Fig. 6) with a time resolution of 500 ms using the 3M™ Bair Hugger™ Temperature Monitoring System. Participants were instructed to avoid having any exercises prior to the experiment.

**Figure 1.**
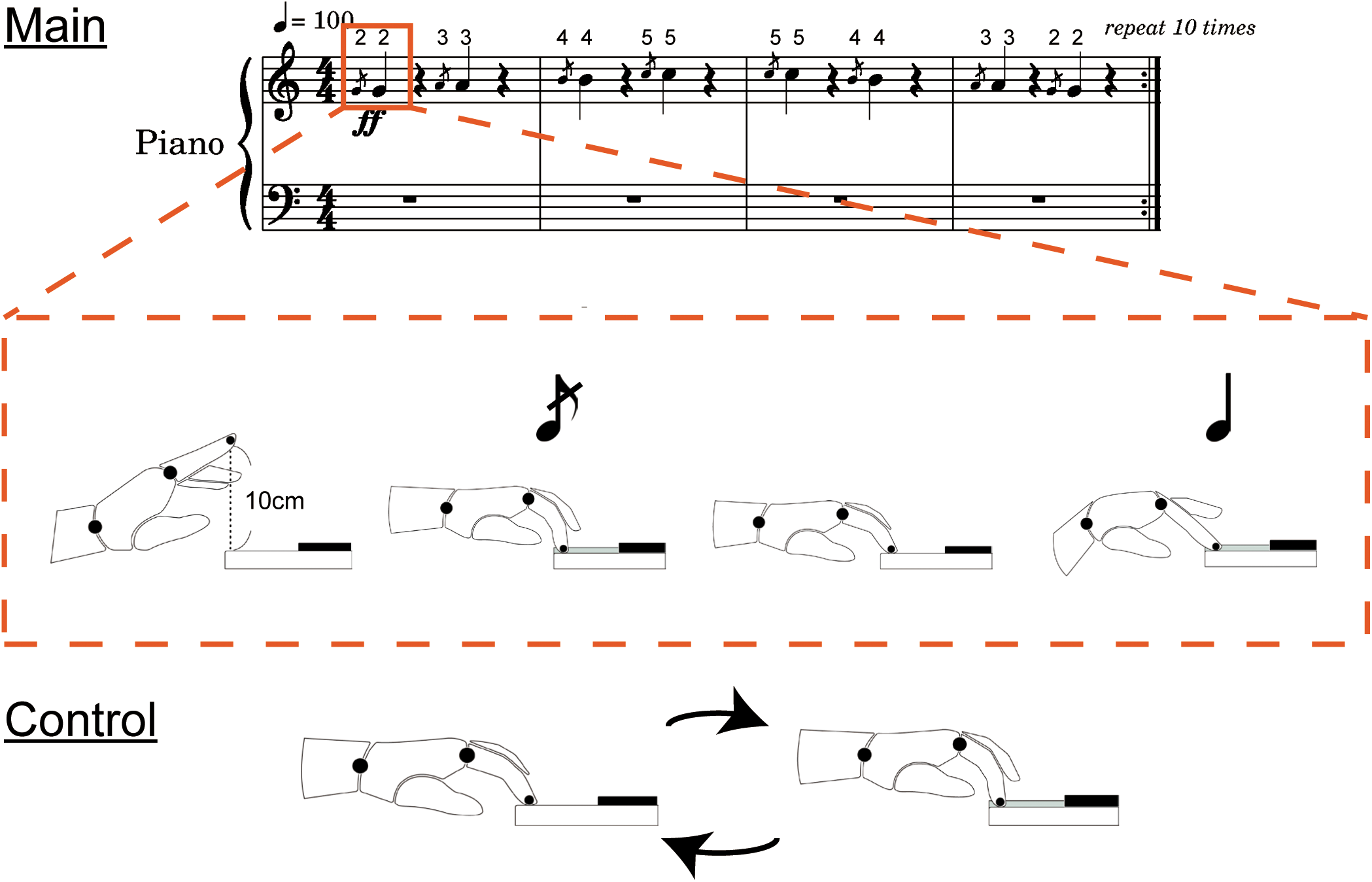
A musical score representing the training task (top panel) and a schematic drawing of the keystroke movements corresponding to the repetitive keystrokes used by the main group with the plyometric training (middle panel) and control group who underwent the same number of keystrokes as the main group (bottom panel). The number on the score represents the fingering (2, 3, 4, and 5 corresponds the index, middle, ring, and little finger, respectively).

**Figure 2.**
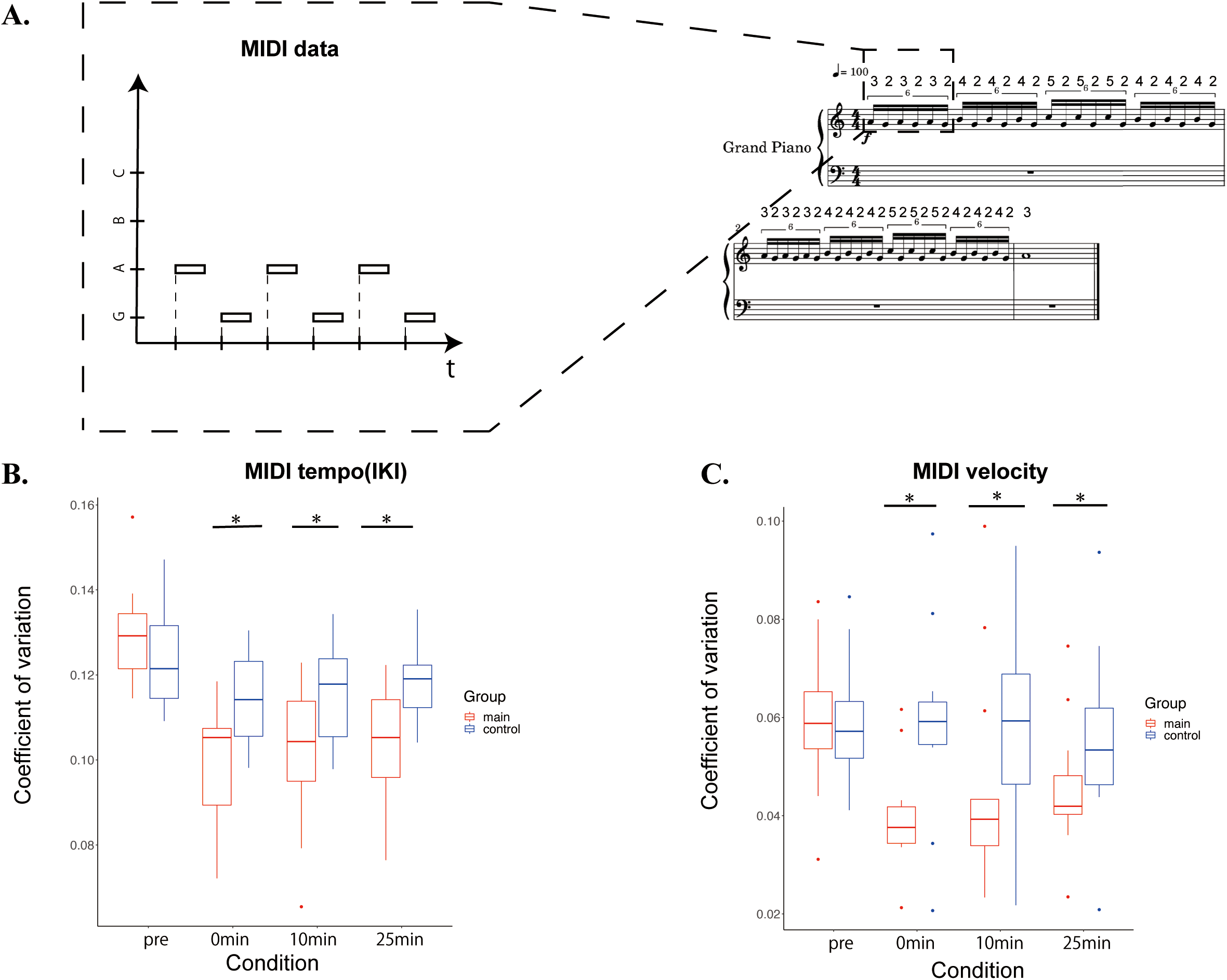
A: A schematic illustration of the temporal information of the individual keystrokes representing a musical score representing the test task. B and C: Box plots of the group means of the coefficients of variation (CV) of the timing (MIDI Inter-keystroke Intervals: IKI) and velocity (MIDI velocity) of the keypresses before and after the training session (i.e. condition in the x-axis) in the main (red box) and control (blue box) groups. *: p<0.05.

**Figure 3.**
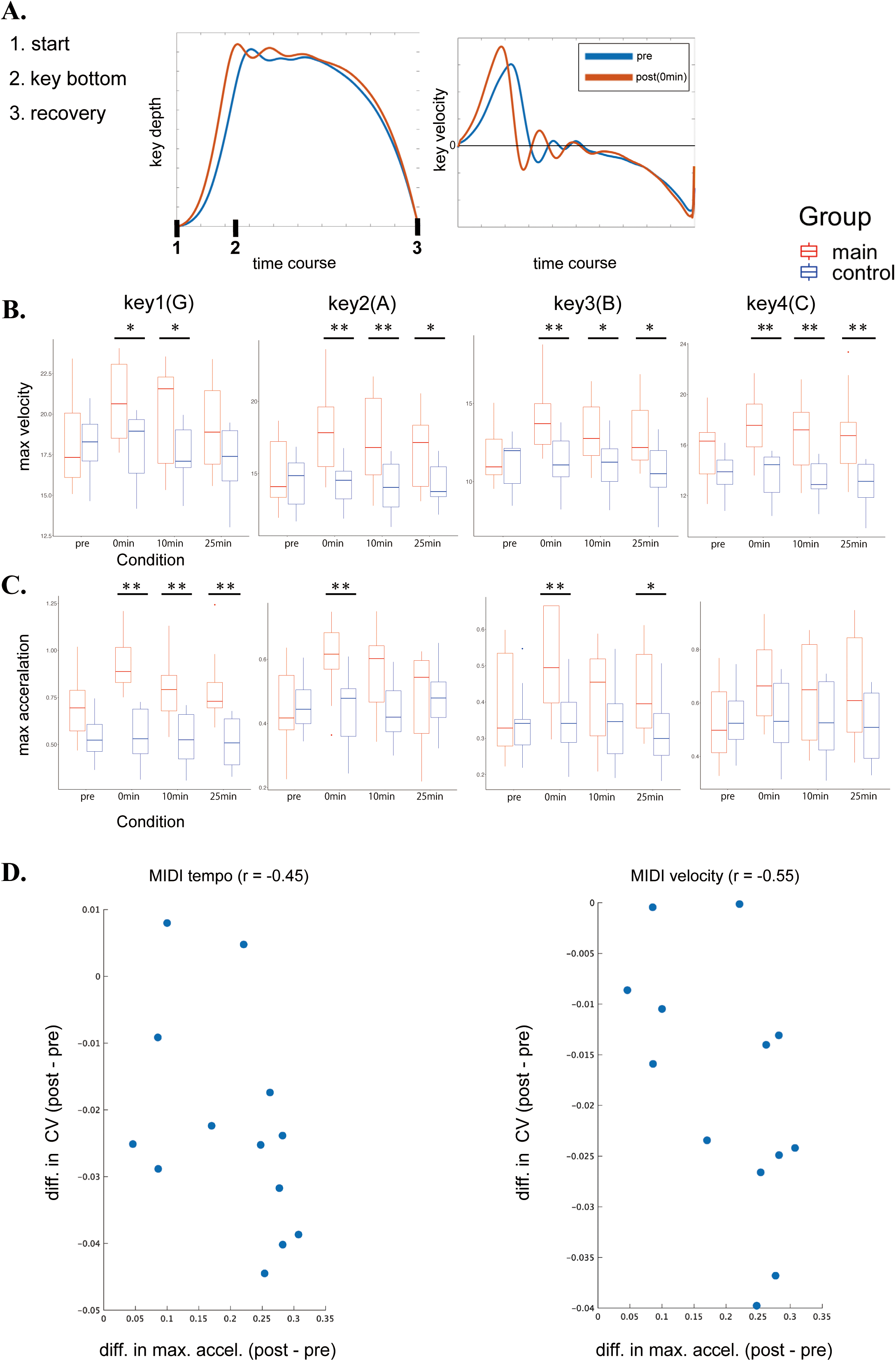
A: Representative examples of the time-varying trajectories of the vertical position of the piano key (left) and their derivatives (right) at the pretest (blue) and posttest (i.e. 0 min after the training) (red) of one representative pianist in the main group. B and C: Box plots of the group means of the maximum descending velocity (B) and acceleration (C) of the trajectories of the four keys to be struck (i.e. key1-4) before and after training (x-axis) in the main (red box) and control (blue box) groups. *p < 0.05, **p<0.01. D: Scatter plots of the differential values between the pretest and posttest in the maximum keystroke acceleration relative to the CV of the keystroke timing (left panel) and velocity (right panel).

**Figure 4.**
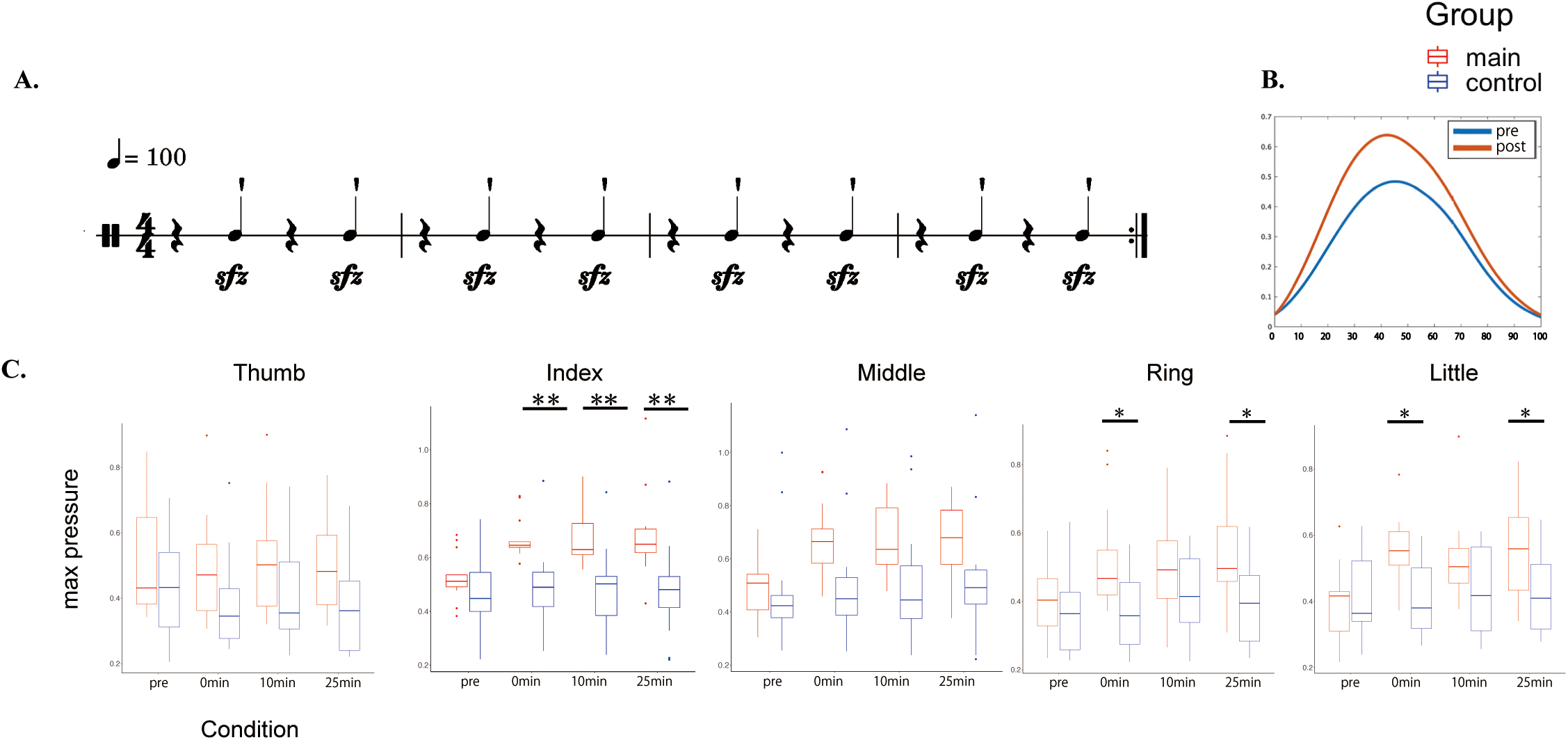
A: A score in the finger force production test. B: Representative trajectories of the finger pressure of one pianist. C: Group means of the maximum pressure exerted by each of the five digits before and after the training task (x-axis; condition) in the main (red) and control (blue) groups. *p < 0.05, **p<0.01.

**Figure 5.**
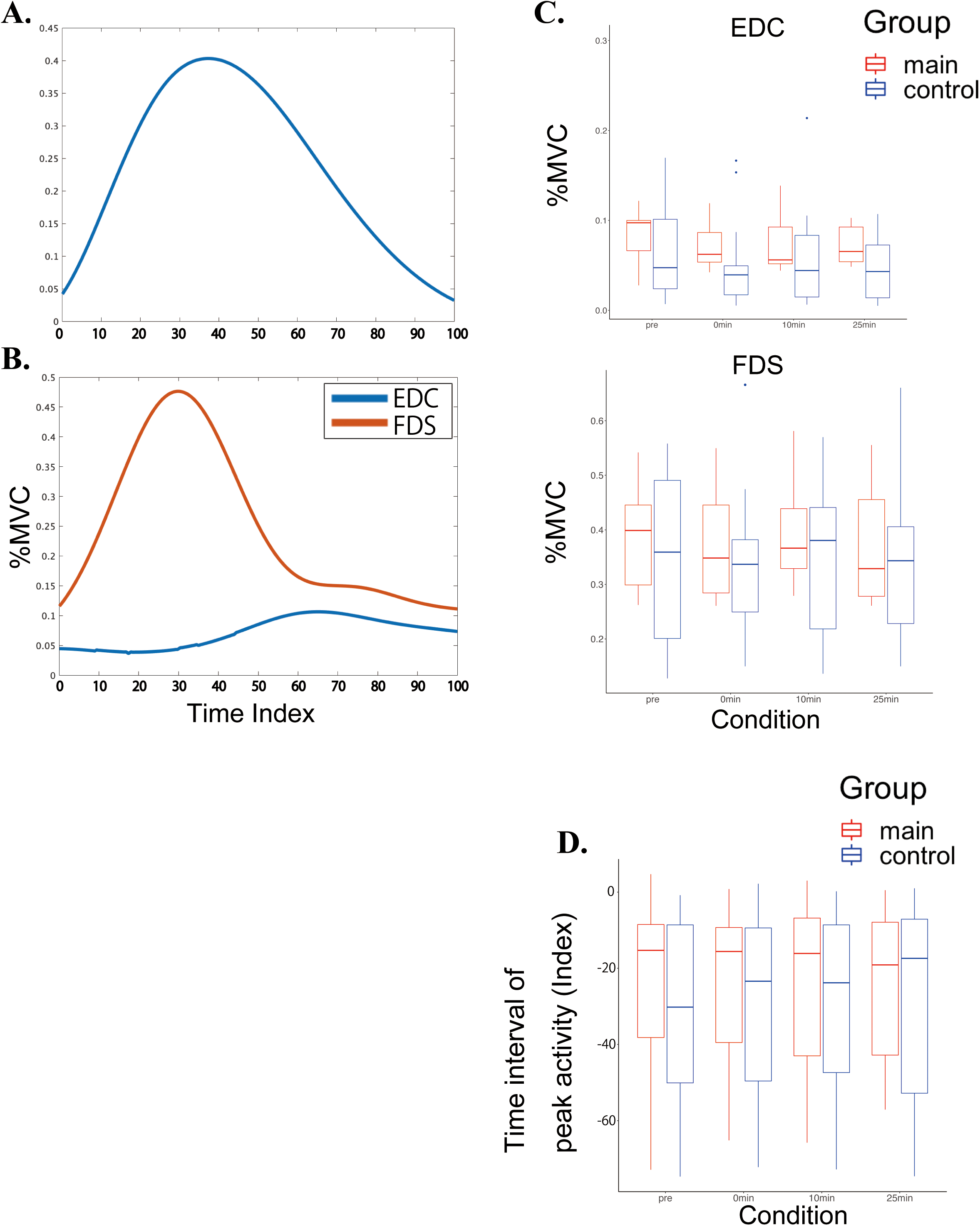
A and B: Representative examples of the time-varying trajectory of the finger pressing force (A) and its corresponding muscular activities of the finger extensor and flexor muscles (i.e. EDC and FDS) (B) of one representative pianist in the main group. The x-axis indicates the normalized time so that the period from the initiation to the termination of the force production can be 100 timepoints. C: Box plots of group means o the maximum values of the muscular activities at the EDC and FDS before and after the training session in the main (red) and control (blue) groups. D: Box plots of group means of the interval of the timing of the peak activities between EDC and FDS in the main and control groups. The negative value indicates when the peak FDS activity preceded the peak EDC activity. *p < 0.05, **p<0.01.

**Figure 6.**
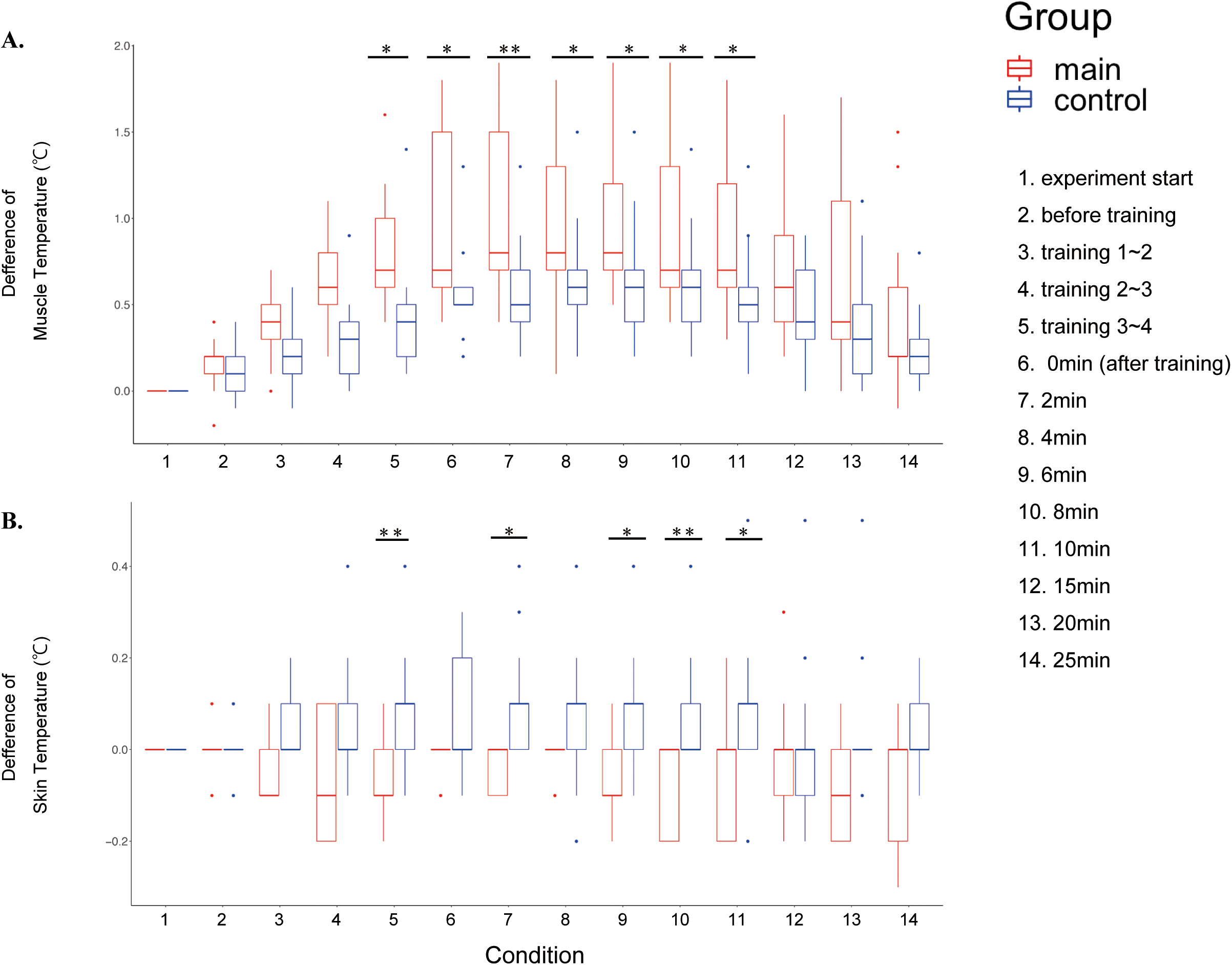
A time-course of the temperature of the finger extensor muscle at the forearm (A) and skin of the forearm (B) throughout a course of the experiment in the main group (with the plyometric training in red) and control group (without the plyometric training in blue). The x-axis indicates the conditions, each of which is indicated at the right box in the figure. *p < 0.05, **p<0.01.

### Experimental Tasks

The experiment consisted of three successive sessions within a single day: pre-test, training, and post-test. In addition, the post-test session consisted of three trials with a break in between; 0min, 10min, and 25min after the training. In the pre-test and post-test sessions, participants were asked to perform two tasks: the piano test and finger force production test. At the beginning of the experiment, the isometric maximal voluntary contraction (MVC) was asked to be performed at the EDC and FDS to calculate %MVC during the task performance.

For the piano test, participants played the melody designated on a test score (see Fig. 2A right) with the piano by the right hand, while their elbow was put on a table to minimize motions of the other body portions (i.e. forearm only). Furthermore, they were instructed so that the fingers could be kept on the surface of the keyboard from the beginning of the keystroke as much as possible and that the wrist could be immobilized without any rotational movements. The participants were asked to play with maintaining the tempo of 100 BPM (i.e. with the inter-tone duration of 100 msec due to sextuplet) as accurately as possible during the task performance with keeping the loudness as consistent as possible. The tempo was provided with a metronome only before each performance was initiated.

For the finger force production test, the elbow and wrist were immobilized on a table, and only the fingers were used to press the force sensor in a manner displayed on the musical score (see Fig. 4A). As in the piano test, their fingertips were always kept contact with the force sensors. Then the participants were asked to press as strongly and quickly as possible isometrically, along with the tempo provided by the metronome.

In the training session, participants were asked to perform the training task with the right hand in an instructed manner that differed between the groups. Participants in the main group were instructed to perform a plyometric-like exercise with the piano 40 times in a manner displayed on the score (see the score in Fig.1), which was characterized as follows: 1) one continuous cycle of swinging the hand down from 100mm above the keyboard toward the key and returning to the original position, 2) two strong strikes in succession as a unit, the first with the downward elbow motion and the second with the wrist flexion (i.e. snapping), 3) being aware of relaxing the muscles except at the moment of each enunciation, 4) making the interval between two successive strikes as short as possible. By contrast, participants in the control group repeated the same exercise as the aforementioned piano test 40 times, so that the total duration of the training session could be the same between the main and control groups.

### Data analysis

#### Movement variables

The MIDI information obtained from the keyboard was used as variables for evaluating the keystroke performance. The coefficient of variation (CV) of the inter-keystroke interval (IKI) of two successive strikes was used as a variable representing stability of the tempo, whereas the CV of the keystroke velocity was used as an index representing the loudness stability.

Data of the finger pressure and key motions were cut for each press/keystroke as epochs according to a threshold (three times of the standard deviation of the signals prior to the task performance), which was used for time normalization of the epochs. These data were averaged across the epochs for smoothing, and the diff function in MATLAB (Mathworks Inc.) was used to compute the first and second derivatives of the vertical position of the key. The maximum value of each waveform was used for the subsequent analyses.

#### EMG preprocessing

The EMG data were bandpass filtered at 10-250 Hz to remove artificial high-frequency noise and movement artifacts. The same time-index was used for time normalization of the EMG signals to temporally align each epoch with the time normalized force and key motion.

### Statistics

A two-way mixed-design ANOVA (independent variables: Group and Condition) or three-way mixed-design ANOVA (independent variables: Group, Condition, and Finger) was run as needed. If Mauchly’s sphericity test was necessary, the Greenhouse-Geisser correction was performed. Post-hoc was performed only in the case of significance with correction for multiple comparisons (p < 0.05).

## RESULTS

### Training effects on variability of the inter-keystroke intervals and keypress velocity

Figure 2 illustrates the group means of the coefficient of variation of the inter-keystroke interval (Fig.2B) and that of the keypress velocity (Fig.2C) at the piano test (Fig.2A) before and after the training session in the main and control groups. For the rhythmic variability of the keystrokes, a two-way mixed-design ANOVA with group and condition yielded both interaction effect (F(3,72)=3.967, p =1.23 × 10^-4^, η^2^=0.053) and main effect of condition (F(3,72) =14.51, p = 1.73 × 10^-7^, η^2^=0.170), but no main effect of group (F(1,24) = 5.19, p = 0.48,η^2^= 0.014). Post-hoc comparison showed group differences only after the training session. For the inter-strike variability of the keypress velocity, both the interaction effect between group and condition (F(3,72) = 8,736, p = 5.13× 10^-5^, η^2^= 0.048) and main effect of condition (F(3,72) = 11.58, p = 2.80× 10^-6^, η^2^= 0.062) were significant, whereas there was no main effect for group (F(1,24) = 3.029, p =0.095, η^2^= 0.098).

### Training effects on the piano key-descending velocity and acceleration

Figure 3 shows the group means of the maximum descending velocity (Fig.3B) and acceleration (Fig.3C) of the key-motion at the piano test (Fig.2A) before and after the training session in the main and control groups. For the maximum key velocity, a two-way mixed-design ANOVA with group and condition revealed a significant interaction effect as well as main effects of both group and condition for all keys (see Table 1). Post-hoc comparison showed group differences only after the training session for all keys. For the maximum acceleration, the interaction effects were evident at all of the four keys to be struck, whereas the main effects of the group at the key-1 and key-3 and the main effects of condition at all keys were significant (see Table 2). Post-hoc comparison yielded group differences only after the training session for all keys except for the key-4. We also found a negative correlation of the differential value of the maximum acceleration of the key descending motion between the pretest and posttest in the main group, both with the variability of the inter-keystroke intervals (r = −0.45) and that of the keypress velocity (r = −0.55), respectively.

**Table 1.**
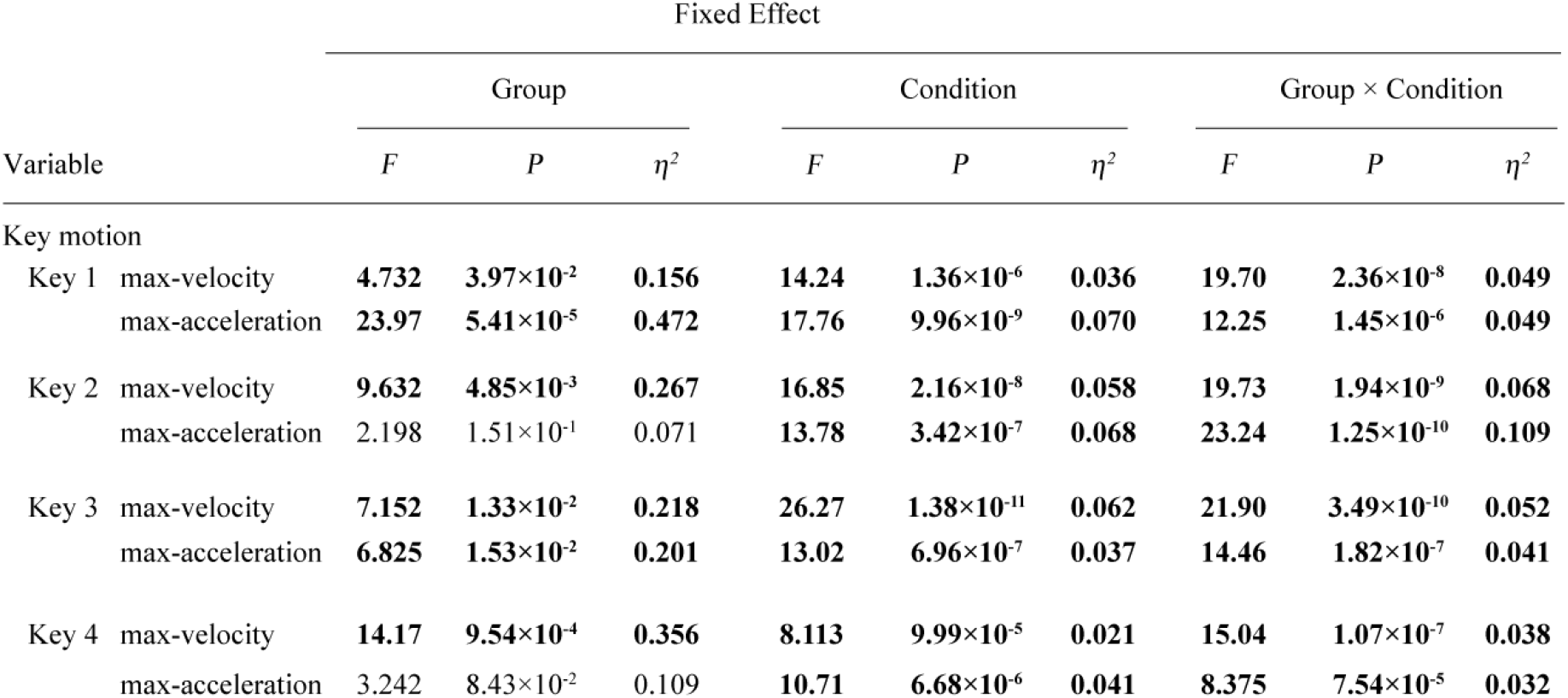
Results of two-way mixed-design ANOVA for the maximum velocity and acceleration of the key-motion

| Variable |  | Fixed Effect |  |  |  |  |  |  |  |  |
| --- | --- | --- | --- | --- | --- | --- | --- | --- | --- | --- |
| | | Group | | | Condition | | | Group × Condition | | |
| | | *F* | *P* | $\eta^2$ | *F* | *P* | $\eta^2$ | *F* | *P* | $\eta^2$ |
| Key motion |  |  |  |  |  |  |  |  |  |  |
| Key 1 | max-velocity | 4.732 | 3.97$\times 10^{-2}$ | 0.156 | 14.24 | 1.36$\times 10^{-6}$ | 0.036 | 19.70 | 2.36$\times 10^{-8}$ | 0.049 |
| | max-acceleration | 23.97 | 5.41$\times 10^{-5}$ | 0.472 | 17.76 | 9.96$\times 10^{-9}$ | 0.070 | 12.25 | 1.45$\times 10^{-6}$ | 0.049 |
| Key 2 | max-velocity | 9.632 | 4.85$\times 10^{-3}$ | 0.267 | 16.85 | 2.16$\times 10^{-8}$ | 0.058 | 19.73 | 1.94$\times 10^{-9}$ | 0.068 |
| | max-acceleration | 2.198 | 1.51$\times 10^{-1}$ | 0.071 | 13.78 | 3.42$\times 10^{-7}$ | 0.068 | 23.24 | 1.25$\times 10^{-10}$ | 0.109 |
| Key 3 | max-velocity | 7.152 | 1.33$\times 10^{-2}$ | 0.218 | 26.27 | 1.38$\times 10^{-11}$ | 0.062 | 21.90 | 3.49$\times 10^{-10}$ | 0.052 |
| | max-acceleration | 6.825 | 1.53$\times 10^{-2}$ | 0.201 | 13.02 | 6.96$\times 10^{-7}$ | 0.037 | 14.46 | 1.82$\times 10^{-7}$ | 0.041 |
| Key 4 | max-velocity | 14.17 | 9.54$\times 10^{-4}$ | 0.356 | 8.113 | 9.99$\times 10^{-5}$ | 0.021 | 15.04 | 1.07$\times 10^{-7}$ | 0.038 |
| | max-acceleration | 3.242 | 8.43$\times 10^{-2}$ | 0.109 | 10.71 | 6.68$\times 10^{-6}$ | 0.041 | 8.375 | 7.54$\times 10^{-5}$ | 0.032 |

**Table 2.** Results of two-way mixed-design ANOVA for each of the finger pressure and activations of the EDC (extensor) and FDS (flexor) muscles in the finger force production task

| Variable | Fixed Effect |  |  |  |  |  |  |  |  |  |  |  |  |  |  |  |  |  |  |  |  |
| --- | --- | --- | --- | --- | --- | --- | --- | --- | --- | --- | --- | --- | --- | --- | --- | --- | --- | --- | --- | --- | --- |
| | Group | | | Condition | | | Finger | | | Group × Condition | | | Group × Finger | | | Condition × Finger | | | Group × Condition × Finger | | |
| | F | P | $\eta^2$ | F | P | $\eta^2$ | F | P | $\eta^2$ | F | P | $\eta^2$ | F | P | $\eta^2$ | F | P | $\eta^2$ | F | P | $\eta^2$ |
| **Finger pressure** |  |  |  |  |  |  |  |  |  |  |  |  |  |  |  |  |  |  |  |  |  |
| Max | 5.660 | 2.56$\times 10^{-2}$ | 0.139 | 25.39 | 4.73$\times 10^{-4}$ | 0.035 | 13.39 | 7.75$\times 10^{-4}$ | 0.109 | 18.91 | 2.10$\times 10^{-2}$ | 0.026 | 1.10 | 3.57$\times 10^{-1}$ | 0.010 | 7.42 | 1.68$\times 10^{-3}$ | 0.019 | 1.850 | 6.01$\times 10^{-2}$ | 0.005 |
| Time to peak | 0.297 | 5.91$\times 10^{-1}$ | 0.006 | 0.272 | 0.781 | 0.002 | 3.702 | 0.0454 | 0.016 | 0.716 | 0.505 | 0.006 | 0.356 | 0.645 | 0.002 | 0.988 | 0.386 | 0.007 | 1.464 | 0.240 | 0.011 |
| **EMG (finger force development task)** |  |  |  |  |  |  |  |  |  |  |  |  |  |  |  |  |  |  |  |  |  |
| Max (EDC) | 0.999 | 1.42$\times 10^{-1}$ | 0.012 | 0.737 | 2.22$\times 10^{-1}$ | 0.006 | 0.626 | 2.70$\times 10^{-1}$ | 0.015 | 0.415 | 7.43$\times 10^{-1}$ | 0.001 | 1.388 | 2.44$\times 10^{-1}$ | 0.002 | 0.711 | 7.40$\times 10^{-1}$ | 0.002 | 0.999 | 4.50$\times 10^{-1}$ | 0.003 |
| Max (FDS) | 2.446 | 1.31$\times 10^{-1}$ | 0.063 | 0.841 | 2.79$\times 10^{-1}$ | 0.004 | 0.948 | 2.69$\times 10^{-1}$ | 0.007 | 1.770 | 8.77$\times 10^{-2}$ | 0.003 | 0.974 | 4.25$\times 10^{-1}$ | 0.009 | 0.677 | 4.43$\times 10^{-1}$ | 0.002 | 1.050 | 3.97$\times 10^{-1}$ | 0.005 |
| Interval of peaks | 1.143 | 2.96$\times 10^{-1}$ | 0.018 | 0.124 | 9.45$\times 10^{-1}$ | 0.0001 | 0.774 | 5.12$\times 10^{-1}$ | 0.036 | 0.755 | 5.23$\times 10^{-1}$ | 0.001 | 2.221 | 9.26$\times 10^{-1}$ | 0.004 | 1.481 | 1.31$\times 10^{-1}$ | 0.007 | 2.883 | 9.91$\times 10^{-1}$ | 0.001 |

### Training effects on the maximum finger force exertion

In order to identify factors associated with the aforementioned results, we investigated effects of training on RFD during the finger force exertion in the finger force production test. Figure 4 shows the group means of the maximum finger force exerted by each of the four fingers (Fig.4C) at the designated finger force production task (Fig.4A) before and after the training in the main and control groups. A three-way mixed-design ANOVA with group, condition, and finger was performed for the maximum exerted force (see Table 2). There was no second-order interaction, whereas significant first-order interactions were found for both Finger × Condition and Group ×Condition, but not for Group × Finger. The main effects of all three factors were also significant.For each finger, post-hoc comparison was conducted for Group × Condition, and overall groupwise differences were evident for the fingers 2, 4, and 5, but not for the fingers 1 and 3. For the time to which the exerted finger force reached its peak value, ANOVA revealed the main effect only of the finger, but none of the interactions nor the other main effects were significant (Table 2).

### Finger muscular activities during the finger force production test

Figure 5 illustrates the group means of the maximum activities of the EDC and FDS muscles (Fig.5C) and the time-varying waveforms of these muscular activities (Fig.5B) along with the force exerted by the index finger (Fig. 5A) during the finger force production test. For the maximum values, a three-way mixed-design ANOVA with group, condition, and finger showed that neither the interactions nor main effects were significant for each of the EDC and FDS (Table 2). Similarly, for the time interval of the peak activities between the EDC and FDS (Fig.5D), a three-way mixed-design ANOVA yielded neither significant interaction nor main effects.

### Training effects on changes of muscle and skin temperature

Figure 6 shows the group means of the time-varying muscle temperatures at the extrinsic finger extensor (EDC) (Fig.6A) and the forearm skin temperatures (Fig.6B) throughout the experiment.A two-way mixed-design ANOVA with condition and group was performed for the muscle temperature and found a significant interaction(F(13,312) = 3.154, p = 4.23×10^-2^, η^2^=0.032), main effects of group (F(1,24) = 5.035, p = 3.43 × 10^-2^, η^2^= 0.136) and condition (F(13,312) = 53.48, p = 6.71× 10^-15^, η^2^= 0.359). Post-hoc comparisons revealed group differences particularly during the period from the second half of the training to 10 min after the training. A two-way mixed-design ANOVA for the skin temperature similarly showed both a significant interaction (F(13,312)=2.602, p= 1.93 × 10^-2^, η^2^=0.049) and main effect of group (F(1,24)=9.968, p =4.26× 10^-3^, η^2^=0.179), but no condition effect (F(13,312)=1.364, p=2.32× 10^-1^, η^2^=0.026). Post-hoc comparisons showed significant group differences during the period similar to that of the muscle temperature.

## DISCUSSION

The present study found that the plyometric training targeting the extrinsic finger flexor muscle was effective as exemplified by a decrease of the variability of both timing and velocity of the keystrokes when performing a pianistic task that requires loud and fast tone production. On the other hand, such a significant effect of the training was not observed following the training with repetitive piano keystrokes that did not involve the plyometric-like muscular contraction (i.e. a control group). The contrasting group difference indicates that the plyometric exercises used in this study enhances precision of the finger movements in fast and forceful repetitive piano keystrokes, which has been difficult to be achieved in previous studies. Interestingly, the spatiotemporal features of the finger muscular activities did not change following the plyometric exercise for both FDS and EDC, whereas the finger muscular but not skin temperature was elevated as the plyometric exercise was being performed. This suggests physiological changes at the finger muscles by the plyometric training. Together, these results indicate that the plyometric exercise has potentials of further improving well-trained performance skills of pianists.

To evaluate the training effect on the finger motor functions, we assessed RFD in the isometric finger force production for flexion. Specifically following the plyometric-like piano training, RFD was increased for each of the four fingers that underwent the training. A lack of any training effect at the thumb that did not perform the plyometric training and at all fingers that underwent the conventional repetitive practicing supports the idea that the enhanced ability of the finger force production resulted from this training. Similarly, the training effect was also evident for the maximum acceleration of the piano key-depression, which was correlated with improvement of precision of timing and velocity in the piano keystrokes. One possible explanation for the enhanced piano performance is a negative physiological relationship between the muscular strength and variability of the force production^27^. It is therefore plausible that the strengthening effect of the plyometric exercise on the finger muscles aided in reducing signal-dependent noise in the motor commands issued into the muscles and thereby decreased the variability of the exerted force. Interestingly, the muscular activation was not augmented through the training, even though the force production was increased. This indicates that the target force can be produced with reduced finger muscular activities of the finger, implicating improvement of physiological efficiency in the finger force production. This can play a role in preventing muscular fatigue and/or development of overuse syndromes through piano practicing, in addition to enhancement of timing and force precision in piano performance.

As one putative physiological mechanism behind the effect of the plyometric training on the force production ability, we found elevation of the finger muscular temperature but not of the skin temperature specifically following this training. Muscle contraction begins with the release of calcium ions from the sarcoplasmic reticulum into the myofibrils, which binds actin and myosin heads (cross-bridges) and then consumes energy from the ATPase reaction for contraction. Previous studies have shown that the cross-bridge cycling rates for muscle contraction are affected largely by the temperature-dependent myosin-ATPase reaction^28–29^, which explains why the muscle temperature was elevated along with the increase in RFD of the skeletal muscles in both present and previous studies ^28,30–31^. In other words, the increase in RFD may be due to an increase in the chemical reaction rate during muscle contraction as the muscle temperature elevates, which has been observed as an increase in the muscle power output in passively warming (water immersion, ~1°C) hands^32^. This can be a potential reason why the RFD was increased through the plyometric training.

The phenomenon of improved motor performance following a conditioning exercise is called PAPE, which is recently proposed to distinguish it from conventional Post Activation Potentiation (PAP) due to Myosin Light Chain Phosphorylation^12^. It has been shown that plyometric exercise can induce PAPE as a conditioning exercise not only in the lower limb but also in the upper limb at the short force production rather than resilience exercises with high and lasting long force production^12,33^. In the present study, we used the plyometric exercise for the finger muscles, as in other studies^34^ advocating PAPE which demonstrated concomitant changes in the finger muscular temperature along with the training, and eventually an increase in the RFD at the four fingers that underwent the training. Several remarkable effect of skill improvement by PAPE has been demonstrated in tasks especially with high-speed force exertion such as sprinting and jumping. Piano performance in this study similarly require high-speed movements and strong muscle contractions of more than 10 times per second, and the results indicate that the application of PAPE is highly compatible with such movements. This observation suggests that selective application of plyometric exercises to the finger extrinsic muscles induced PAPE, which may underlie the improved motor skills in piano performance. Several alternative explanations for the skill improvement still remain, which include that motor learning with a modified rhythm of the target task optimized muscle coordination of performance as was observed in a previous study^9^, or that participants acquired a different way of keystrokes toward approaching the key surface through the training, despite instructions to keep the fingers contact with the key throughout the performance.

One limitation to infer the physiological mechanism of the present training effect is no assessment of chemical changes within the muscles through the training and neuroplastic changes in the primary motor cortex and spinal cord by means of non-invasive brain stimulation techniques. In future studies, it is also necessary to record activities of the intrinsic hand muscles and the other extrinsic muscles in order to uncover the entire physiological mechanism of the plyometric training.

## ACKNOWLEDGEMENT

The present study was supported by JST CREST (JPMJCR20D4), JST CREST (JPMJCR17A3), and JSPS Grant-in-Aid for Transformative Research Areas B (20H05713).

